# Evaluating Specimen Quality and Results from a Community-Wide, Home-Based Respiratory Surveillance Study

**DOI:** 10.1101/2020.12.07.415653

**Authors:** Ashley E. Kim, Elisabeth Brandstetter, Naomi Wilcox, Jessica Heimonen, Chelsey Graham, Peter D. Han, Lea M. Starita, Denise J. McCulloch, Amanda M. Casto, Deborah A. Nickerson, Margaret M. Van de Loo, Jennifer Mooney, Misja Ilcisin, Kairsten A. Fay, Jover Lee, Thomas R. Sibley, Victoria Lyon, Rachel E. Geyer, Matthew Thompson, Barry R. Lutz, Mark J. Rieder, Trevor Bedford, Michael Boeckh, Janet A. Englund, Helen Y. Chu, on behalf of the Seattle Flu Study Investigators

## Abstract

**Introduction:** While influenza and other respiratory pathogens cause significant morbidity and mortality, the community-based burden of these infections remains incompletely understood. The development of novel methods to detect respiratory infections is essential for mitigating epidemics and developing pandemic-preparedness infrastructure.

**Methods:** From October 2019 to March 2020, we conducted a home-based cross-sectional study in the greater Seattle area, utilizing electronic consent and data collection instruments. Participants received nasal swab collection kits via rapid delivery within 24 hours of self-reporting respiratory symptoms. Samples were returned to the laboratory and were screened for 26 respiratory pathogens and a human marker. Participant data were recorded via online survey at the time of sample collection and one week later.

**Results:** Of the 4,572 consented participants, 4,359 (95.3%) received a home swab kit, and 3,648 (83.7%) returned a nasal specimen for respiratory pathogen screening. The 3,638 testable samples had a mean RNase P C_R_T value of 19.0 (SD: 3.4) and 1,232 (33.9%) samples had positive results for one or more pathogens, including 645 (17.7%) influenza-positive specimens. Among the testable samples, the median time between shipment of the home swab kit and completion of laboratory testing was 8 days [IQR: 7.0-14.0].

**Discussion:** Home-based surveillance using online participant enrollment and specimen self-collection is a feasible method for community-level monitoring of influenza and other respiratory pathogens, which can readily be adapted for use during pandemics.

## Introduction

Acute respiratory illnesses (ARIs) constitute a significant burden on the healthcare system in the United States and represent an important cause of morbidity and mortality worldwide [1-4]. In the United States, influenza causes 140,000 - 810,000 hospitalizations and 12,000 - 67,000 deaths annually [1-4]. Additionally, respiratory syncytial virus (RSV) leads to approximately 2 million outpatient visits each year for children under the age of 5 [5,6]. Estimates of the prevalence of ARI-causing pathogens generally rely on in-person healthcare visits or aggregate counts from hospitalized individuals [6-10]. Thus, these estimates likely omit cases of mild to moderate ARI in community-dwelling individuals who may not seek care for their illness [11-13].

Active, community-level monitoring of respiratory infections is essential to assess the seasonal activity of ARI-causing pathogens and can be used to inform public health prevention strategies and influence treatment decisions made at the community level. Previous respiratory pathogen surveillance studies evaluated specific subsets of the population, such as households with children, or used labor-intensive, coordinated efforts to capture a representative sample of the community, which makes such approaches difficult to replicate [14-16]. Additionally, similar to traditional respiratory surveillance networks, some of these studies relied on healthcare facility visits which have the potential to result in the nosocomial spread of respiratory pathogens [17-18]. Despite the limitations of earlier analyses, community-wide surveillance studies remain of vital importance as they provide opportunities to better understand the epidemiology of respiratory illness among symptomatic individuals with variable disease severities and healthcare-seeking behaviors.

The Seattle Flu Study Swab and Send sub-study is a novel, city-wide, cross-sectional study of home-based detection of respiratory pathogens. This study demonstrates the feasibility of using a home-based surveillance approach to assess the epidemiology of influenza and other respiratory pathogens in a community-based setting.

## Methods

### Study Design

The “Swab and Send” sub-study was nested within the Seattle Flu Study (SFS), a multi-armed influenza surveillance system [19]. This sub-study aimed to assess the feasibility of city-wide home-based cross-sectional respiratory pathogen surveillance, utilizing rapid delivery systems for at home collection of a nasal swab from individuals experiencing ARIs with return of specimens to the laboratory for respiratory pathogen detection. Individuals residing within the greater Seattle area with ARI symptoms were prospectively enrolled from October 2019 - March 2020. Participants resided in 89 different zip codes within King County in and around the city of Seattle. This study was approved by the University of Washington Institutional Review Board.

### Recruitment

Study recruitment occurred through 1) referrals from healthcare providers, clinics, Seattle Flu Study community kiosks (an in-person enrollment center), schools, and workplaces, 2) dissemination of printed flyers posted at community locations, and 3) posting of targeted online advertisements (e.g., Facebook, Instagram, Twitter, Google). Recruitment materials directed potential participants to the study website (www.seattleflu.org, henceforth referenced as the “study website”). To determine their eligibility, individuals completed a screening survey on the study website by providing their age, home zip code, and information about the presence and duration of respiratory symptoms and by verifying their access to the internet.

Individuals were eligible to participate in the study if they lived within specified zip codes, had experienced new or worsening cough and/or two ARI symptoms (subjective fever, headache, sore throat or itchy/scratchy throat, nausea or vomiting, runny/stuffy nose or sneezing, fatigue, muscle or body aches, increased trouble with breathing, diarrhea, ear pain/ discharge, or rash) within seven days of enrollment (Table A1), were English-speaking, had a valid email address, and had access to the internet at home. All individuals consented to participate in the research study electronically, with consent by a parent or legally-authorized representative for individuals under 18 years and concurrent assent for those between 7 and 18 years.

### Data Collection

Upon consenting, participants completed an online *Enrollment Questionnaire* to provide their home address and contact information such as an email address or phone number. Participants were mailed a home swab kit within 48 hours of submitting the *Enrollment Questionnaire*, which included a *Quick Start Instruction Card* (Fig. A1), a universal viral transport media (UTM) tube (Becton, Dickinson and Company, Sparks, MD), a nylon flocked mid-turbinate swab (COPAN Diagnostics Inc., Murietta, CA), a return box with an affixed Category B UN3373 label (as required by International Air Transport Association (IATA) guidelines [20]), and a pre-paid return shipping label. Pediatric nasal swabs (COPAN Diagnostics Inc., Murietta, CA) were available for participants 5 years of age or younger. Various couriers were used to deliver home swab kits to participants across King County, depending on geographical location as determined by zip code. For the 2,398 of participants who resided within the city of Seattle, FedEx Same Day City was used to deliver kits with a target delivery time of two hours.

Upon kit receipt, participants completed an online *Illness Questionnaire* to ascertain demographics, illness characteristics, and health behaviors. Education level was only asked to participants 18 and older. Additionally, participants were asked to rate the impact of their current illness on regular activities at the time of their enrollment using a five-point Likert scale with the following levels: not at all, a little bit, somewhat, quite a bit, or very much. These categories were transformed into none, low (a little bit, somewhat), and high (quite a bit, very much).

At the end of the *Illness Questionnaire*, participants were prompted to self-collect a mid-nasal swab using the provided *Quick Start Instruction Card* (Fig. S1) included in the swab kit box. Participants were instructed to place their self-collected nasal swabs directly into the UTM tube which was pre-labeled with a unique sample barcode. Next, participants were instructed to place the UTM tube containing the self-collected nasal swab into a specimen bag, pre-packaged with an absorbent sheet, and then to put the specimen bag into the provided return shipping box. United States Postal Service (USPS) return postage and Category B UN3373 stickers were affixed to the outside of the return box. Although previous testing has demonstrated that respiratory viral RNA is stable at room-temperature in UTM for up to one week [21], participants were encouraged to return their nasal specimen within 24 hours or as soon as possible. For the subset of participants where detailed courier data was available, median delivery times were determined through the use of proof of delivery (POD) data on scheduled shipment times, completed delivery times, and mileage.

Seven days after nasal swab collection, participants were re-contacted to complete a *One Week Follow-Up Questionnaire* to assess the impact of their illness on behavioral outcomes such as absenteeism and healthcare-seeking behaviors (provider visits, antiviral use, etc). Care-seeking was marked as “any care” if the participant indicated they had sought care in the *Illness Questionnaire* or *One Week Follow-Up Questionnaire*. Any care-seeking included doctor’s office or urgent care, pharmacy, hospital or emergency department, or other.

All study questionnaires were collected through REDCap (Table A3) [22]. A full timeline of study events may be found in Table A2. Access to de-identified, aggregate study data and analysis code will be publicly available on the study website.

### Laboratory Testing

When kits arrived in the study laboratory, contents of the box and deviations from return mail instructions were recorded. 200 µl of UTM was removed and subjected to RNA extraction using a MagNA Pure 96 System (Roche) and the remainder was banked at −80°C. The extracted nucleic acids were screened for respiratory pathogens using a custom, TaqMan-based Open Array panel (Thermo Fisher) and an additional SARS-CoV-2 RT-PCR research assay. Samples were subjected to the SARS-CoV-2 assay in real-time if they were collected after February 25, 2020 and retrospectively if collected between January 1, 2020 and February 24, 2020 (Table A4) [23]. Samples with RNase P relative cycle threshold (C_RT_) values ≤28 for the Open Array assay, which has a preamplification step, and ≤36 for the SARS-CoV-2 assay were considered to contain sufficient material for pathogen detection [24]. Samples were screened for influenza A H3N2, H1N1, and pan influenza A, influenza B, influenza C, respiratory syncytial viruses (RSV) A and B, human coronaviruses (hCoV) 229E, NL63, OC43, and HKU1, SARS-CoV-2, adenovirus (AdV), human rhinovirus (hRV), human metapneumovirus (hMPV), human parechovirus (hPeV), enteroviruses A, B, C, D, D68, and G, human bocavirus (hBoV), *Streptococcus pneumoniae, Mycoplasma pneumoniae*, and *Chlamydia pneumoniae* (Table A4). C_RT_ values for RNase P, influenza, hCoV, RSV, and hRV from 11,984 nasal samples collected between October 2019 to March 2020 at Seattle Children’s Hospital were analyzed as a contemporary control of healthcare worker-collected specimens and compared to the self-collected specimens in this study.

### Data Analyses

Descriptive statistics were performed for categorical and continuous covariates. Bivariate analyses were conducted using parametric and nonparametric tests as appropriate, with statistical significance defined as p<0.05. The Kruskal-Wallis test was used to determine p-values for study procedure compliance categories, comparing each of the three nasal swab error types to those with no errors. ANOVA was used to calculate an overall p-value for RNase P values across confidence and discomfort levels. Respiratory pathogen prevalence is defined as the total number of cases detected out of the total number of tested samples.

## Results

### Participant Characteristics

A total of 4,572 participants were consented and enrolled in the SFS Swab and Send sub-study from October 16, 2019 to March 9, 2020. The majority of participants were recruited into the study through online or social media advertisements (53.9%) or through referrals from friends or family (19.3%). Of the 4,572 participants who completed the electronic consent form, 4,359 (95.3%) participants also completed the *Enrollment Questionnaire* and provided a valid home address, which was required to receive a home swab kit. Participant characteristics, including age, sex, race, Hispanic ethnicity, income, education level, influenza vaccination status, healthcare-seeking status, test results, baseline impact of illness on regular activities, and recruitment method are shown in Table 1. The mean age of study participants was 36.6 (SD: 15) years old. Most (73.7%) of participants were 18-49 years old. On average, the study population was more highly educated and had a higher household income than the general population of King County. A total of 31.4% of participants had a bachelor’s degree as their highest degree while 31.6% had an advanced degree. 26.6% had a household income of ≥ $150,000 per year (Table 1).

**Table 1:** Clinical and sociodemographic characteristics of enrolled participants, October 16, 2019 - March 9, 2020

|  |  |
| --- | --- |
|  | N=4,359 (%) |
| **Age** |  |
| <5y | 128 (2.9%) |
| 5-17y | 208 (4.8%) |
| 18-49y | 3212 (73.7%) |
| 50-64y | 614 (14.1%) |
| >=65y | 192 (4.4%) |
| **Sex** |  |
| Male | 1191 (27.3%) |
| Female | 2451 (56.2%) |
| Other | 19 (0.4%) |
| **Race** |  |
| American Indian/ Alaska Native | 17 (0.4%) |
| Asian | 724 (16.6%) |
| Native Hawaiian/ Pacific Islander | 7 (0.2%) |
| Black/African American | 37 (0.8%) |
| White | 2542 (58.3%) |
| Other | 92 (2.1%) |
| Multiple | 188 (4.3%) |
| **Hispanic ethnicity (N=2856)** | 183 (4.2%) |
| **Income** |  |
| ≤\$25K | 196 (4.5%) |
| \$25-50K | 367 (8.4%) |
| \$50-100K | 860 (19.7%) |
| \$100-150K | 738 (16.9%) |
| ≥\$150K | 1160 (26.6%) |
| **Education level** |  |
| Graduated high school/obtained GED or less | 109 (2.5%) |
| Some college (including vocational training, associate's degree) | 492 (11.3%) |
| Bachelor's degree | 1371 (31.5%) |
| Advanced degree | 1377 (31.6%) |
| **Care-seeking** |  |
| Any care prior to enrollment or during study period | 1182 (27.1%) |
| No care prior to enrollment or during study period | 2183 (50.1%) |
| **Illness impact on regular activities at enrollment** |  |
| None | 243 (5.6%) |
| Low | 1597 (36.6%) |
| High | 1831 (42.0%) |
| **How participant heard about the study** |  |
| Saw an ad on Facebook/Instagram/Twitter | 1369 (31.4%) |
| Referral from a friend/family member | 841 (19.3%) |
| Other online | 667 (15.3%) |
| Saw an ad on Google | 314 (7.2%) |
| Referral from my place of work | 280 (6.4%) |
| Other | 172 (3.9%) |
| Saw a Seattle Flu Study kiosk | 86 (2.0%) |
| Email/Seattle Community Pulse | 86 (2.0%) |
| Referral from a healthcare provider, travel clinic, or immigrant/refugee health screening | 60 (1.4%) |
| Referral from my child's school | 29 (0.7%) |

At time of enrollment, 42.0% of participants who were sent a nasal swab rated the impact of their current illness on their regular activities as high although 67.5% had not sought clinical care. The majority of study participants did not seek clinical care for their illness during the study period. A total of 27.1% of participants sought clinical care for their current illness prior to enrollment or during the study period whereas 50.1% never sought clinical care during this time frame (Table 1). In general, participants who sought care were more likely to do so after enrolling and completing their home swab kits. Among those who sought care (N=1,178), 727 (61.7%) participants sought care prior to enrollment and 989 (84.0%) sought care within one week after enrollment, though these categories are not mutually exclusive.

Of the 4,359 participants who received a home swab kit, 3,648 (83.7%) returned a nasal specimen to the laboratory and 3,638 (99.7%) of returned specimens contained sufficient UTM in the tube and RNase P levels for respiratory pathogen screening (Fig. 1). Influenza A (10.8%), hRV (10.4%), hCoV (8.6%), and influenza B (6.9%) were the most commonly detected pathogens (Table A5; Fig. 2). Samples collected on or after January 1, 2020 were tested for SARS-CoV-2, of which 36 out of 2,843 (1.2%) were positive for the novel coronavirus. The 3,629 self-collected nasal specimens with available RNase P data yielded a mean RNase P C_RT_ value of 19.0 (SD: 3.4) (Table A5). A contemporary comparison of C_RT_ values from healthcare worker-collected nasal specimens to self-collected nasal specimens is shown in Table A6.

**Figure 1:**
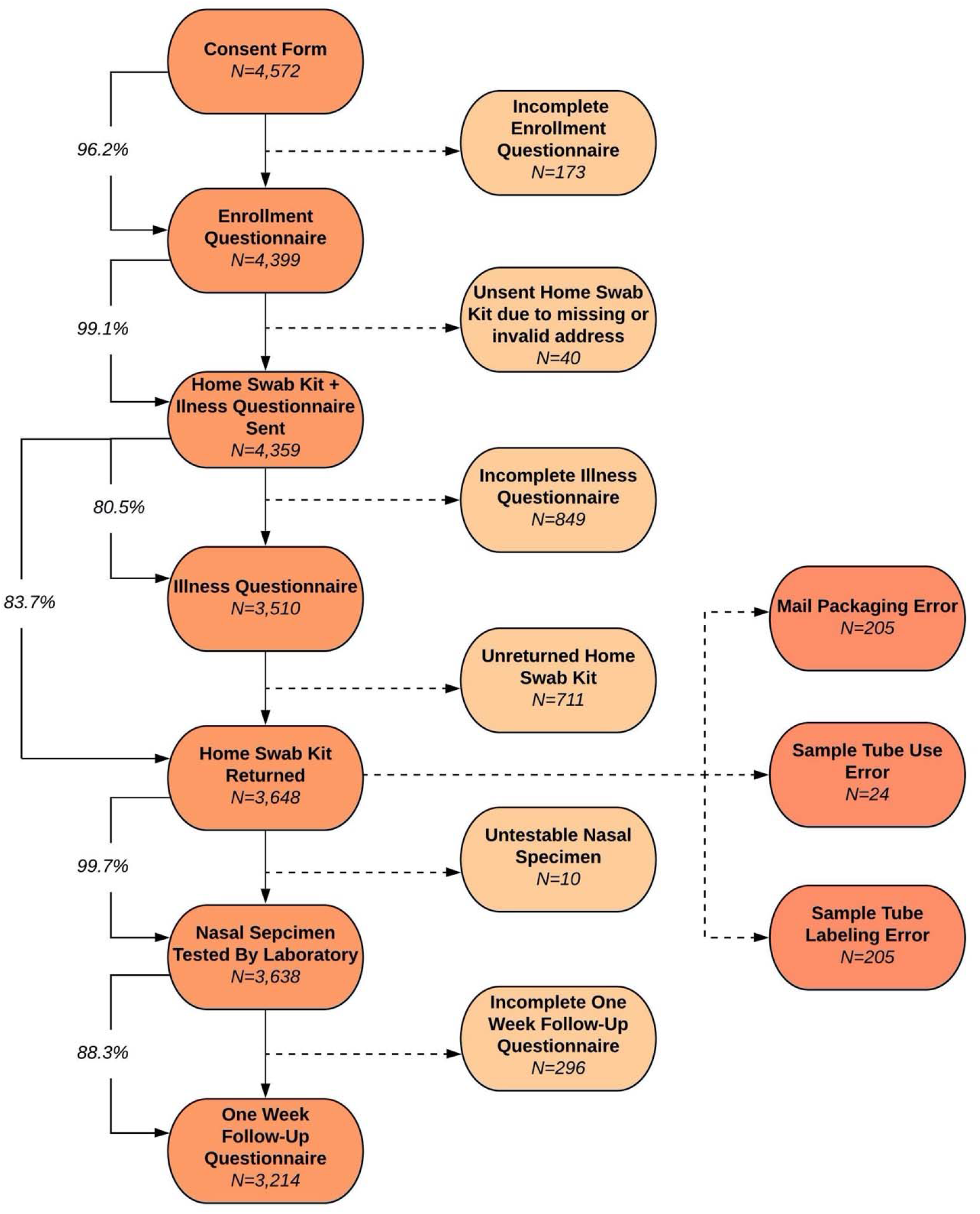
Study procedure completion rates

**Figure 2:**
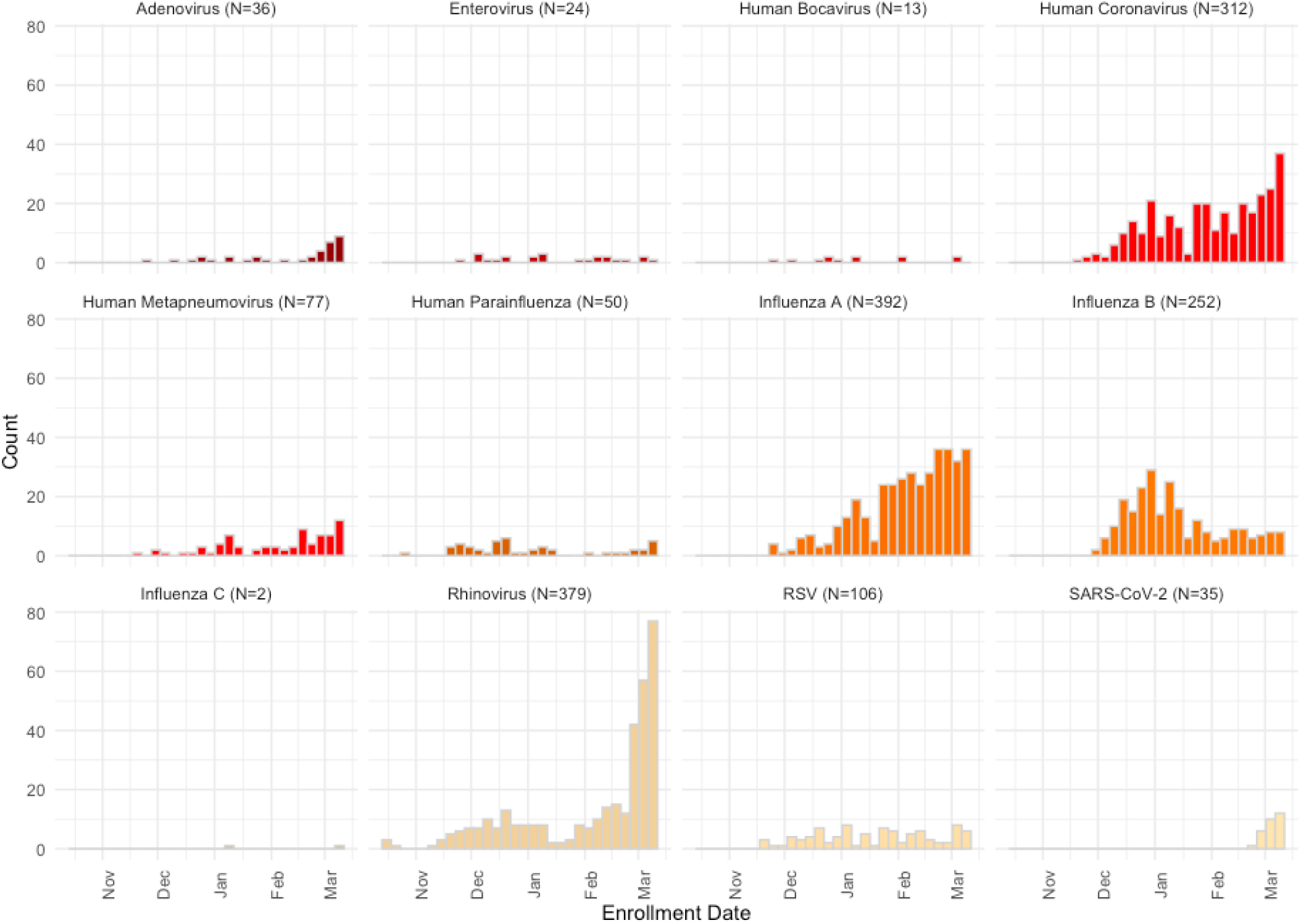
Pathogens detected in participants over time

### Study Logistics

For the 4,359 participants who received a home swab kit, the median time between participant completion of enrollment and scheduling of the shipment was 7.2 hours [IQR: 0.45-19.6]. The total median delivery transit time to participants who received their home swab kit via FedEx Same Day City was 2.2 [IQR: 1.7 - 3.0] hours with 79% of deliveries meeting the two-hour target delivery time. A subset of the delivery time data was reported previously [25]. The median delivery time via FedEx Same Day City to participants’ homes by distance from the study laboratory is shown in Fig. 3. Of the 2,398 FedEx Same Day City deliveries, there were a total of 78 (3.3%) redelivery attempts. The estimated median time between nasal swab collection to receipt at the study laboratory was 3.0 [IQR: 2.0, 4.0] days for the 3,648 participants who returned specimens. Of the 3,638 testable samples, the median time between shipment and completed laboratory testing was 8.0 [IQR: 7.0 - 14.0] days (Table 2).

**Table 2:** Study Logistics & Turnaround Time Metrics

| Metric | Time (Median [IQR]) |
| --- | --- |
| Completed enrollment to shipment scheduled (hours)<sup>a</sup> | 7.2 [0.45-19.6] |
| Delivery time to participant's home (hours)<sup>b</sup> | 2.2 [1.7-3.0] |
| Nasal swab collection to returned specimen received at the laboratory (days)<sup>c</sup> | 3.0 [2.0-4.0] |
| Shipment of home swab kit to participant's home to completed laboratory testing (days)<sup>d</sup> | 8.0 [7.0-14.0] |

<sup>a</sup> Time between participant completion of the *Enrollment Questionnaire* and scheduled shipment of the home swab kit
<sup>b</sup> FedEx Same Day City was used to rapidly deliver home swab kits within the city of Seattle. Median FedEx Same Day City delivery times from ordering the shipment to arrival at the participant's residence, adjusting for redeliveries.
<sup>c</sup> Estimated time between nasal swab collection, measured by completion of the *Illness Questionnaire & Nasal Swab Collection* survey, and the return of completed home swab kits at the laboratory. Completed home swab kits were returned via pre-paid USPS Priority Mail.
<sup>d</sup> Time between scheduled shipment of home swab kits to participants' homes and available results for tested specimens

**Figure 3:**
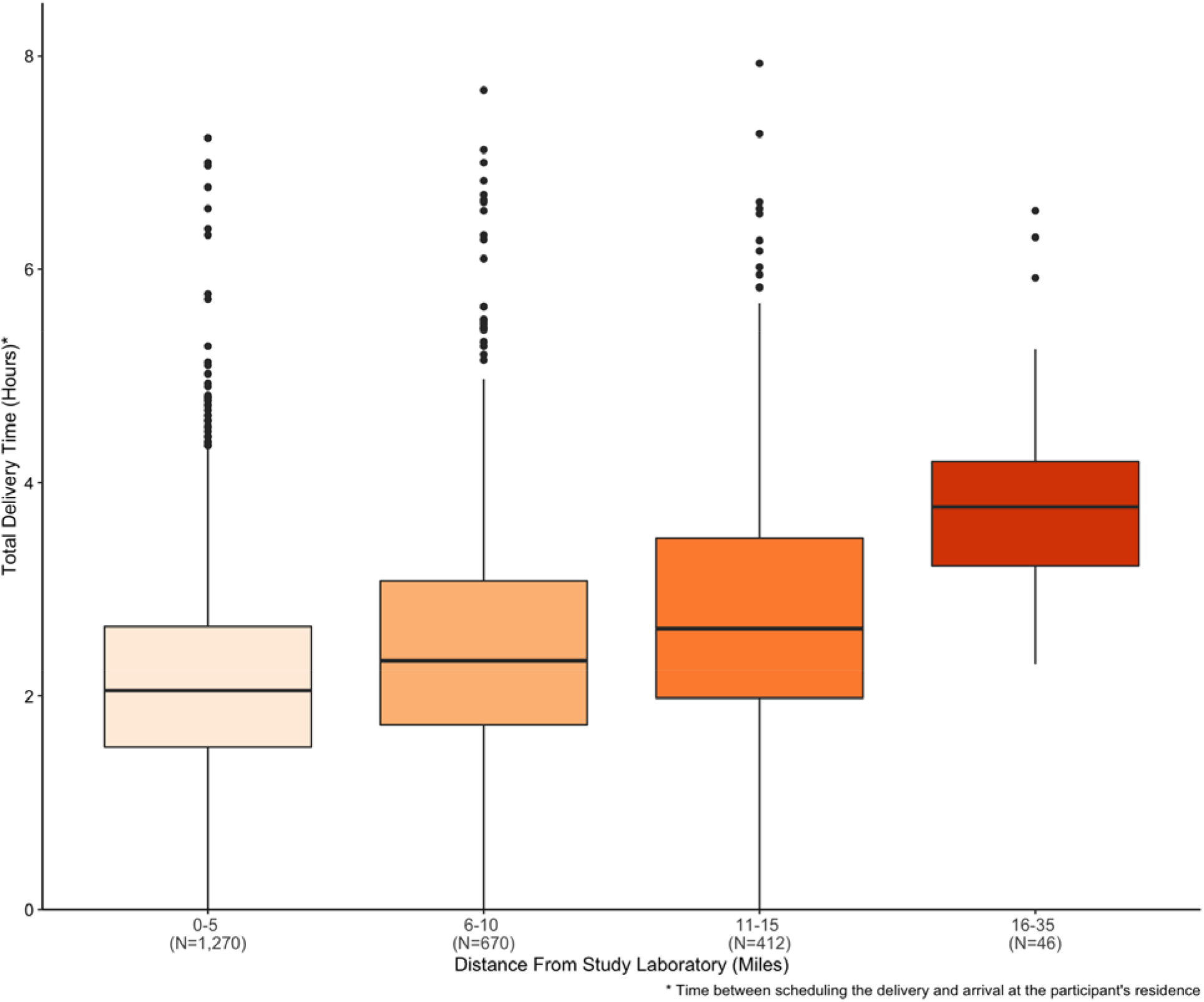
Median delivery times of home swab kits to participants by distance from study laboratory (N=2,398)

### Study Procedure Completion and Compliance

Study procedure completion rates are shown in Fig. 1. Of the 4,359 participants who completed the *Enrollment Questionnaire* and received a home swab kit, 3,214 (73.9%) completed all study procedures. Study procedure completion and compliance by age, sex, income, education, care-seeking status, and baseline illness-impact are shown in Table 3. None of these variables were significantly associated with study procedure compliance (Table 3).

**Table 3:** Clinical and sociodemographic characteristics of enrolled participants, October 16, 2019 - March 9, 2020 by study procedure completion and compliance

|  | Study procedure completion |  | Study procedure compliance |  |  |  |  |
| --- | --- | --- | --- | --- | --- | --- | --- |
|  | Returned Nasal Swab<br>N=3638 | Completed All Study Procedures<br>N=3214 | Mail Packaging Error<sup>†</sup><br>N=205 | Sample Tube Use Error<sup>§</sup><br>N=24 | Sample Tube Labeling Error<sup>¶</sup><br>N=205 | No Packaging or Sample Tube Errors<br>N=3211 | P value* |
| **Age** |  |  |  |  |  |  | 0.11 |
| <5y | 110 (3.0%) | 89 (2.8%) | 9 (4.4%) | 1 (4.2%) | 6 (2.9%) | 92 (2.9%) |  |
| 5-17y | 173 (4.8%) | 149 (4.6%) | 12 (5.9%) | 0 (0%) | 16 (7.8%) | 149 (4.6%) |  |
| 18-49y | 2638 (72.5%) | 2324 (72.3%) | 144 (70.2%) | 15 (62.5%) | 141 (68.8%) | 2339 (72.8%) |  |
| 50-64y | 545 (15.0%) | 496 (15.4%) | 33 (16.1%) | 6 (25.0%) | 29 (14.1%) | 480 (14.9%) |  |
| >=65y | 168 (4.6%) | 153 (4.8%) | 6 (2.9%) | 2 (8.3%) | 10 (4.9%) | 150 (4.7%) |  |
| **Sex** |  |  |  |  |  |  | 0.38 |
| Male | 1142 (31.4%) | 1013 (31.5%) | 70 (34.1%) | 8 (33.3%) | 70 (34.1%) | 994 (31.0%) |  |
| Female | 2340 (64.3%) | 2178 (67.8%) | 115 (56.1%) | 13 (54.2%) | 118 (57.6%) | 2097 (65.3%) |  |
| Other | 18 (0.5%) | 15 (0.5%) | 3 (1.5%) | 0 (0%) | 1 (0.5%) | 14 (0.4%) |  |
| **Income** |  |  |  |  |  |  | 0.81 |
| <= \$25K | 180 (4.9%) | 161 (5.0%) | 5 (2.5%) | 1 (4.2%) | 9 (4.4%) | 164 (5.1%) | |
| \$25-50K | 344 (9.5%) | 315 (9.8%) | 21 (10.3%) | 2 (8.3%) | 27 (13.2%) | 294 (9.2%) | |
| \$50-100K | 818 (22.5%) | 760 (23.6%) | 39 (19.0%) | 10 (41.7%) | 49 (23.9%) | 716 (22.3%) | |
| \$100-150K | 700 (19.2%) | 639 (19.9%) | 33 (16.1%) | 2 (8.3%) | 32 (15.6%) | 635 (19.8%) | |
| >=\$150K | 1129 (31.0%) | 1042 (32.4%) | 69 (33.7%) | 6 (25.0%) | 48 (23.4%) | 1010 (31.5%) | |
| **Education level** |  |  |  |  |  |  | 0.53 |
| Graduated high school/obtained GED or less | 101 (2.8%) | 80 (2.5%) | 9 (4.4%) | 0 (0%) | 10 (4.9%) | 81 (2.5%) |  |
| Some college (including vocational training, associate's degree) | 449 (12.3%) | 414 (12.9%) | 20 (9.8%) | 5 (20.8%) | 32 (15.6%) | 395 (12.3%) |  |
| Bachelor's degree | 1324 (36.4%) | 1220 (38.0%) | 67 (32.7%) | 5 (20.8%) | 58 (28.3%) | 1189 (37.0%) |  |
| Advanced degree | 1328 (36.5%) | 1229 (38.2%) | 68 (33.2%) | 10 (41.7%) | 66 (32.2%) | 1188 (37.0%) |  |
| **Care-seeking** |  |  |  |  |  |  | 0.80 |
| Any care prior to enrollment or during study period | 1138 (31.3%) | 1077 (33.5%) | 52 (25.4%) | 7 (29.2%) | 63 (30.7%) | 1013 (31.5%) |  |
| No care prior to enrollment or during study period | 2136 (58.7%) | 2136 (66.5%) | 114 (55.6%) | 13 (54.2%) | 105 (51.2%) | 1912 (59.5%) |  |
| **Illness impact on regular activities at enrollment** |  |  |  |  |  |  | 0.07 |
| None | 234 (6.4%) | 203 (6.3%) | 17 (8.3%) | 1 (4.2%) | 11 (5.4%) | 205 (6.4%) |  |
| Low | 1521 (41.8%) | 1373 (42.7%) | 90 (43.9%) | 10 (41.7%) | 73 (35.6%) | 1345 (41.9%) |  |
| High | 1754 (48.2%) | 1637 (50.9%) | 81 (39.5%) | 10 (41.7%) | 107 (52.2%) | 1564 (48.7%) |  |

\* Kruskal-Wallis test used to determine p-values for study procedure compliance categories (excludes
first three columns)
† Mail packaging errors include returning the nasal specimen in a damaged box, a different box than the
one provided, an improperly closed box, or an improperly used specimen transport bag or lack thereof
§ Sample tube use errors include returned nasal specimens with a damaged or broken UTM tube, an
absent swab, or leakage
¶ Sample tube labeling errors include a missing written full name or date of collection on the UTM tube

The majority of participants correctly followed instructions to package their collected nasal swab for return to the laboratory. Of the 3,648 returned nasal specimens, 3,208 (88.1%) home swab kits were returned correctly packaged. A total of 205 (5.6%) contained a sample tube labeling error, such as a missing written name or collection date, and 205 (5.6%) were mispackaged. Criteria for mispackaged samples included improper use of the provided return box, specimen transport bag, or lack thereof. Additionally, 24 (0.66%) returned specimens had a sample tube use error, such as a damaged UTM tube, a missing or misused nasal swab, or leakage. Four out of 3,648 (0.11%) returned home swab kits contained leakage and these samples were immediately disposed of upon unpackaging (Table 3).

Participants who enrolled between January 6, 2020 and March, 9, 2020 were asked to rate their confidence in correctly self-collecting their nasal swab and their discomfort level while doing so. Higher confidence and discomfort levels were significantly associated with lower RNase P C_RT_ values (p<0.001 and p=0.04, respectively). The average RNase P C_RT_ value for participants who experienced strong discomfort was 1.4 lower than the average value for those who had no discomfort. The average RNase P C_RT_ value for those who were very confident was 1.2 lower than those who were not confident at all (Fig. 4). Among the 4,359 participants who received a home swab kit, there was one (0.0%) reported adverse event related to strong discomfort while collecting the nasal swab. The affected participant’s discomfort resolved within two minutes. The participant suffered no long-term effects and did not require medical attention. Results suggest that non-medically trained individuals can safely and adequately collect a nasal sample from themselves or their family members.

**Figure 4:**
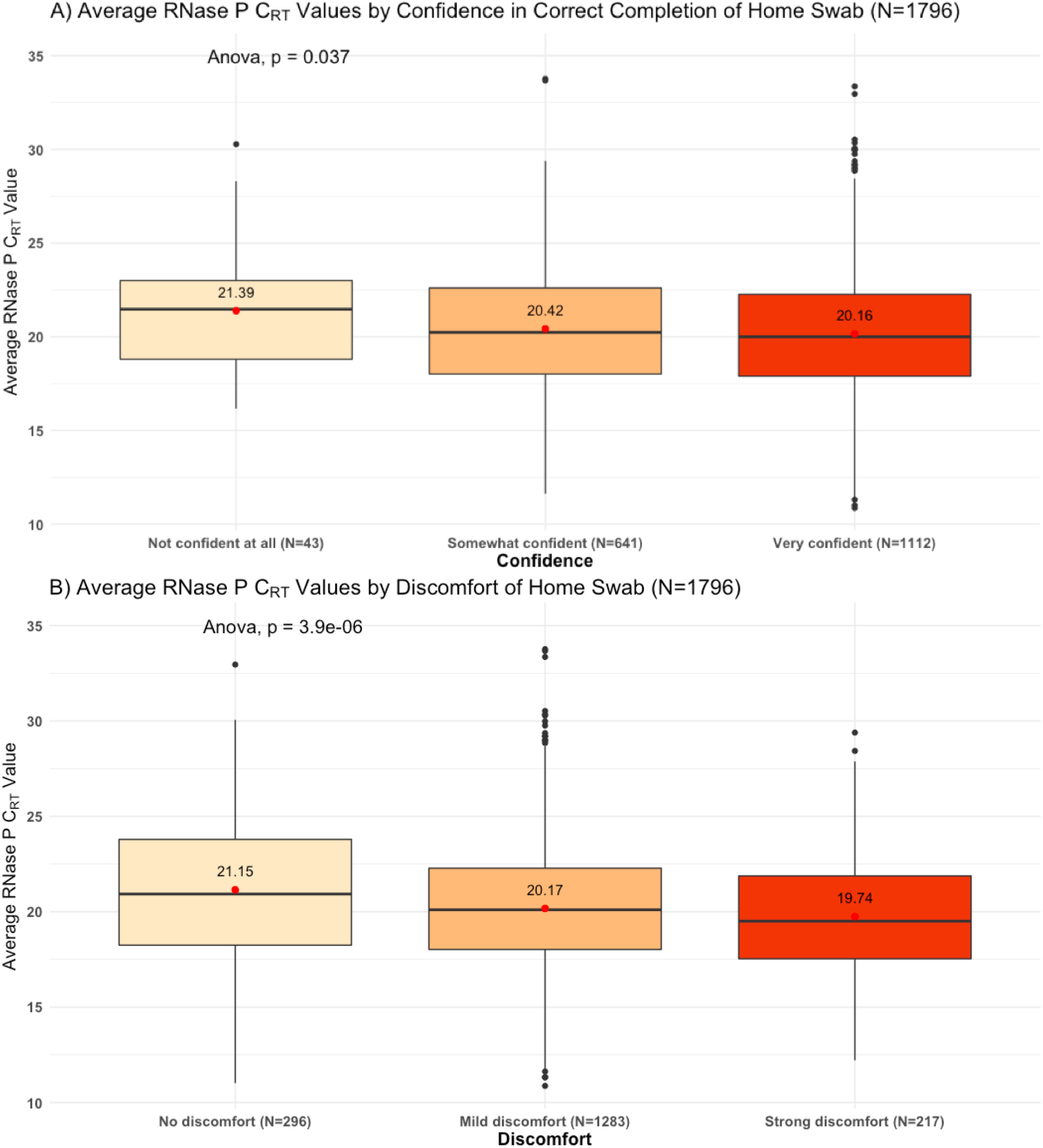
Average RNase P C_RT_ values by discomfort of and confidence in home swab collection

## Discussion

Over the 2019-2020 influenza season, we enrolled a large cohort of participants with acute respiratory illness in a study of home-based swab collection for detection of respiratory pathogens. The majority of participants completed all study procedures and returned their nasal specimens to the study laboratory in a timely manner and in compliance with federal transport guidelines for biohazards. The majority of returned nasal specimens were adequately self-collected as quantified by RNase P C_RT_ value. These results support the feasibility of using online enrollment and self-collected nasal swabs for community surveillance of respiratory pathogens.

Existing methods to estimate the community-level prevalence of influenza rely on estimator models based on laboratory-confirmed cases and adjusted for various confounding factors including medical care seeking, collection and testing of specimens, and reporting of cases. These methods are limited to medically attended illnesses and require relatively comprehensive data for accuracy, which leads to long periods of time between data collection and the availability of results [19]. In this study, we directly surveyed for influenza and other respiratory pathogens in the community allowing rapid assessment of pathogen characteristics and the associated clinical presentations among both care-seeking and non-care-seeking study populations. When combined with estimator models, on-the-ground surveillance of community-dwelling individuals with less severe illness and a wider range of demographic backgrounds may enhance our understanding of the burden of various respiratory pathogens in a community.

Similarly, estimator models with complete reliance on laboratory-confirmed cases can be limiting, especially during epidemics or pandemics in heavily-affected regions where outbreak dynamics are rapidly evolving and the capacity of the healthcare system to adequately test cases has been exceeded [19]. The benefits of direct, home-based surveillance among community-dwelling individuals can be seen in context of the current COVID-19 pandemic. From January 1, 2020 to March 9, 2020, the Seattle Flu Study detected 78 cases of SARS-CoV-2 through direct sampling of community members including the first documented case of community transmission in the US, with 36 cases identified through the Swab and Send sub-study [25, 26]. This study enrolled and tested a large cohort of individuals with ARI symptoms across a large geographical area, half of whom did not seek clinical care prior to or during the study period. The at home study design proved to be an effective means of studying individuals infected with influenza and other respiratory pathogens, many of whom may not have been captured by traditional clinic or hospital surveillance. This demonstrates that when faced with an emerging infectious disease, home-based testing can identify cases among non-care-seeking individuals, providing essential information for pandemic identification, spread, and management.

Limitations of this study include the enrollment of a study population that was not representative of the greater Seattle area. King County demographic data from the 2010 census shows that 49.8% of residents were male and 21.4% were 17 years of age and under, whereas our study population included 27.3% males and 7.7% minors. Additionally, the King County population is 6.0% black or African American and 8.9% Hispanic individuals whereas our study cohort was only 0.8% black or African American and 4.2% Hispanic. The median King County household income in 2016 was $78,800 per year whereas the largest proportion (26.6%) of participants had a household income of greater than $150,000 per year [27]. We hypothesize that factors related to lack of internet access and unfamiliarity with online systems may have contributed to lack of representativeness among certain groups in our study population. The utilization of targeted recruitment strategies aimed at enrolling a larger proportion of participants who were underrepresented in this cohort including males, children, minorities, and individuals of lower socioeconomic statuses could be implemented to yield a more representative study population.

Additionally, while most participants returned their home swab kits with no packaging or sample tube use errors at a rate concordant with a previous study [28], improvements to instructions (e.g. inclusion of instructional videos) may decrease these error rates. Further limitations of this study include use of self-collected mid-nasal swabs, which are not the gold standard for respiratory pathogen detection. However, our group has previously demonstrated that self-collected mid-nasal swabs are highly concordant with health care worker-collected nasopharyngeal swabs for detection of SARS-CoV-2 [29], with results comparable to those of previous studies on the detection of viral pathogens by patient-collected mid-nasal swabs [30-33]. In addition, the contemporary control analysis included in this study shows that C_RT_ values for pathogen-positive samples collected by healthcare workers are comparable to those of self-collected samples, with C_RT_ values for healthcare-collected swabs lower for some targets but higher for others than self-collected swabs. Finally, the requirement of internet access and delivery addresses that are easily accessible by standard shipping couriers may limit the scalability of this method in low resource or rural settings.

In conclusion, at home surveillance with self-collected nasal swabs is a feasible method to study the community-based prevalence of influenza during seasonal epidemics on a city-wide scale. This methodology can be adapted to study a variety of respiratory pathogens affecting diverse study populations with the ability to scale-up to larger sample sizes. In particular, this approach allows for the inclusion of non-care-seeking individuals in respiratory pathogen surveillance studies and may be especially useful during epidemics or pandemics when quarantine and social distancing measures are in place to reduce transmission risks.

## Acknowledgements

The Seattle Flu Study is funded by Gates Ventures. The funder was not involved in the design of the study, does not have any ownership over the management and conduct of the study, the data, or the rights to publish.

Helen Y. Chu, Janet A. Englund, Michael Boeckh, Mark J. Rieder, Matthew Thompson, Barry R. Lutz, Deborah A. Nickerson, Lea M. Starita, and Trevor Bedford designed the study including the laboratory and data informatics procedures. Ashley E. Kim wrote the manuscript, developed the data collection instruments and logistics infrastructure for study implementation, and managed day-to-day responsibilities of the study. Naomi Wilcox performed the data analysis for the manuscript. Chelsey Graham developed the logistics infrastructure of the study. Elisabeth Brandstetter managed the IRB and assisted with quality assurance of the study. Denise J. McCulloch contributed to the implementation and quality assurance of the study. Jessica Heimonen wrote the background section of the manuscript and critically revised the manuscript. Victoria Lyon and Rachel E. Geyer contributed to the design of the home-collection kits, including the Quickstart Instructions Card, as well as managing the kit fabrication procedures. Peter D. Han managed laboratory procedures of the study. Misja Ilcisin, Kairsten A. Fay, Jover Lee, and Thomas R. Sibley contributed to the databasing, informatics, and data preparation of the study. Margaret M. Van de Loo and Jennifer Mooney contributed to the recruitment procedures of the study. Amanda M. Casto helped to edit the manuscript.

We would also like to acknowledge Lincoln Pothan, Mariah Anyakora, Grace Kim, and Miguel Martinez for their assistance in the day-to-day shipping responsibilities for the study, Sarah Sohlberg for assisting with participant communication and overall study support, Jack Henry Kotnik, Kara De Leon, Angel Wong, Rose Marzan, Eshin Ang, Regina Garvey, Peiyu Yi, Ashley Bender, Ashley Song, and Kendall Escene for their role in home swab kit fabrication, and Audrey Obsterbind for her support in study implementation.

## Seattle Flu Study Investigators

### Principal Investigators

Helen Y. Chu, MD, MPH^1,7^, Michael Boeckh, MD, PhD^1,2,7^, Janet A. Englund, MD^3,7^, Michael Famulare, PhD^4^, Barry R. Lutz, PhD^5,7^, Deborah A. Nickerson, PhD^6,7^, Mark J. Rieder, PhD^7^, Lea M. Starita, PhD^6,7^, Matthew Thompson, MBChB, MPH, DPhil^9^, and Jay Shendure, MD, PhD^6,7,8^ and Trevor Bedford, PhD^2,6,7^

### Co-Investigators

Amanda Adler, MS^3^, Elisabeth Brandstetter, MPH^1^, Roy Burstein, PhD^4^, Amanda M. Casto, MD, PhD^1,2^; Shari Cho, MS^7^, Anne Emanuels, MPH^1^, Chris D. Frazar, MS^6^, Rachel E. Geyer, MPH^9^, Peter D. Han, MS^7^, James Hadfield, PhD^1^, Jessica Heimonen, MPH^1^, Michael L. Jackson, PhD, MPH^10^, Anahita Kiavand, MS^7^, Ashley E. Kim, BS^1^, Louise E. Kimball, PhD^2^, Jack Henry Kotnik, BA^9^, Kirsten Lacombe, RN, MSN^3^, Jennifer K. Logue, BS^1^, Victoria Lyon, MPH^9^, Denise McCulloch, MD, MPH^1^, Jessica O’Hanlon, BS^1^, Matthew Richardson BA^6^, Julia Rogers, MPH^1^, Thomas R. Sibley, BA^2^, Monica L. Zigman Suchsland, MPH^9^, Melissa Truong, BS^7^, Caitlin R. Wolf, BS^1^ and Weizhi Zhong, BS^7^.

### Affiliations

1. Department of Medicine, University of Washington
2. Vaccine and Infectious Disease Division, Fred Hutchinson Cancer Research Center
3. Seattle Children’s Research Institute
4. Institute for Disease Modeling
5. Department of Bioengineering, University of Washington
6. Department of Genome Sciences, University of Washington
7. Brotman Baty Institute
8. Howard Hughes Medical Institute
9. Department of Family Medicine, University of Washington
10. Kaiser Permanente Washington Health Research Institute

