## Supplementary Material for "Evaluating Specimen Quality and Results from a Community-Wide, Home-Based Respiratory Surveillance Study"

**Appendixes**

**Table A1.** List of symptoms that were used in online questionnaires to screen individuals for eligibility. Selecting either acute cough *or* two or more concurrent *qualifying* symptoms were considered an acute illness episode and made an individual eligible for enrollment in the Swab and Send study.

| Feeling feverish or warm * | Runny/stuffy nose or sneezing * |
| --- | --- |
| Headache * | Feeling more tired than usual * |
| New or worsening cough ** | Muscle or body aches * |
| Chills or shivering ^X^ | Increased trouble with breathing * |
| Sweats ^X^ | Diarrhea ^+^ |
| Sore throat or itchy/scratchy throat * | Ear pain/ear discharge ^+^ |
| Nausea or vomiting * | Rash ^+^ |

* A qualifying symptom for study eligibility for individuals of any age

** A qualifying symptom that is sufficient on its own for study eligibility for individuals of any age

^X^ Not a qualifying symptom for study eligibility

^+^ A qualifying symptom for study eligibility for individuals <18 years of age

**Table A2:** Timeline of events for each participant

|  | **Study Day -2** | **Within 48 hours of enrollment** | **Study Day 0** | **Study Day 7** | **Within 1 month of Study Day 7** |
| --- | --- | --- | --- | --- | --- |
| e-Consent | X |  |  |  |  |
| *Enrollment Questionnaire* | X |  |  |  |  |
| Swab kit received by participant |  | X |  |  |  |
| *Illness Questionnaire* |  |  | X |  |  |
| Self-collect nasal swab and return to the laboratory via a pre-paid return shipping label |  |  | X |  |  |
| *One Week Follow-Up Questionnaire* |  |  |  | X |  |
| Receive electronic gift card claim code after completion of One Week Follow-up Questionnaire |  |  |  |  | X |

**Table A3:** REDCap Survey Instruments

| **Instrument Name** | **Trigger for instrument** | **Who Completes** |
| --- | --- | --- |
| Eligibility Screening (Seattle Flu Study website enrollment wizard) | Seattleflu.org connects with REDCap | Participant |
| e-Consent for Swab and Send | Seattleflu.org sorts individuals into “Swab and Send” study arm | Participant/Legally authorized representative |
| *Enrollment Questionnaire* | e-Consent completed | Participant |
| *Illness Questionnaire & Nasal Swab Collection* | Study staff ships a swab kit to the participant’s house | Participant |
| *Post Collection Data Entry and Quality Control* | Upon arrival at the University of Washington BBI Laboratory | Lab Staff |
| *One Week Follow-Up Questionnaire* | Automatically generated email containing a survey link will be sent to the participant seven days after completion of the *Illness Questionnaire* | Participant |

**Table A4.** Pathogens for which all Seattle Flu Study respiratory specimens are tested using a TaqMan RT-PCR.

| **Viruses** | **Bacteria** |
| --- | --- |
| SARS-CoV-2^1^ | *Streptococcus pneumoniae* |
| Influenza A - H3N2 | *Mycoplasma pneumoniae* |
| Influenza A - H1N1 | *Chlamydia pneumoniae* |
| Influenza A - Pan |  |
| Influenza B |  |
| Influenza C |  |
| Respiratory syncytial viruses A and B |  |
| Parainfluenza viruses 1-4 |  |
| Coronaviruses 229E, NL63, OC43, and HKU1 |  |
| Adenovirus |  |
| Rhinovirus |  |
| Human metapneumovirus |  |
| Human parechovirus |  |
| Enterovirus^2^ |  |
| Enterovirus D68 |  |
| Human Bocavirus |  |

^1^ SARS-CoV-2 was tested for using a stand-alone assay whereas the remaining pathogens were tested for using the Open Array assay

^2^ All enterovirus species A, B, C, D, and G, including: all Coxsackie serotypes under species A, B, C; all Echovirus serotypes; and all Poliovirus serotypes (1-3).

**Table A5:** Virological characteristics of enrolled participants, October 16, 2019-March 9, 2020

|  | **Total (%)** |
| --- | --- |
| **Any positive test result (N=3,509)** | 1232 (33.9%) |
| **Test result* (N=3,638)** |  |
| Influenza A | 392 (10.8%) |
| Influenza B | 252 (6.9%) |
| Influenza C | 2 (0.1%) |
| RSV | 106 (2.9%) |
| SARS-CoV-2 (N=2,843)** | 36 (1.0%) |
| hRV | 379 (10.4%) |
| PIV (I-IV) | 50 (1.4%) |
| hCoV | 312 (8.6%) |
| hBoV | 13 (0.4%) |
| AdV | 36 (1.0%) |
| hMPV | 77 (2.1%) |
| Enterovirus | 24 (0.7%) |
| **Coinfection (N=3,638)*** | 74 (2.0%) |
| **RNase P C_RT_ value (N=3,629), Mean (SD)** | 19.0 (3.4) |

* Results are not mutually exclusive

** Note: Only samples collected on or after January 1, 2020 were tested for SARS-CoV-2.

**Table A6**: Contemporary control comparison of healthcare worker-collected to self-collected nasal specimens from October 2019 to March 2020

| **Seattle Children’s Hospital (N=11,984)** | **RNase P (N=4,463)** | **Influenza (N=2,660)** | **RSV (N=856)** | **hCoV (N=580)** | **hRV (N=2,366)** |
| --- | --- | --- | --- | --- | --- |
| Mean (SD) | 13.4 (2.6) | 15.0 (6.0) | 17.7 (8.0) | 20.2 (6.8) | 24.7 (7.2) |
| Median [Min, Max] | 13.3 [5.6, 30.8] | 13.8 [1.8, 24.9] | 16.5 [1.8, 38.2] | 19.7 [3.4, 39.8] | 25.6 [2.3, 39.8] |
| **Swab and Send (N=3,637)** | **RNase P (N=3,629)** | **Influenza (N=644)** | **RSV (N=106)** | **hCoV (N=312)** | **hRV (N=379)** |
| Mean (SD) | 19.0 (3.4) | 18.7 (4.9) | 18.4 (5.1) | 18.1 (5.1) | 20.9 (4.6) |
| Median [Min, Max] | 18.7 [9.6, 33.8] | 18.9 [5.2, 27.7] | 18.5 [17.8, 39.2] | 17.8 [6.8, 27.7] | 21.0 [8.14, 27.8] |

**Fig. A1:** *Quick Start Instruction Guide*

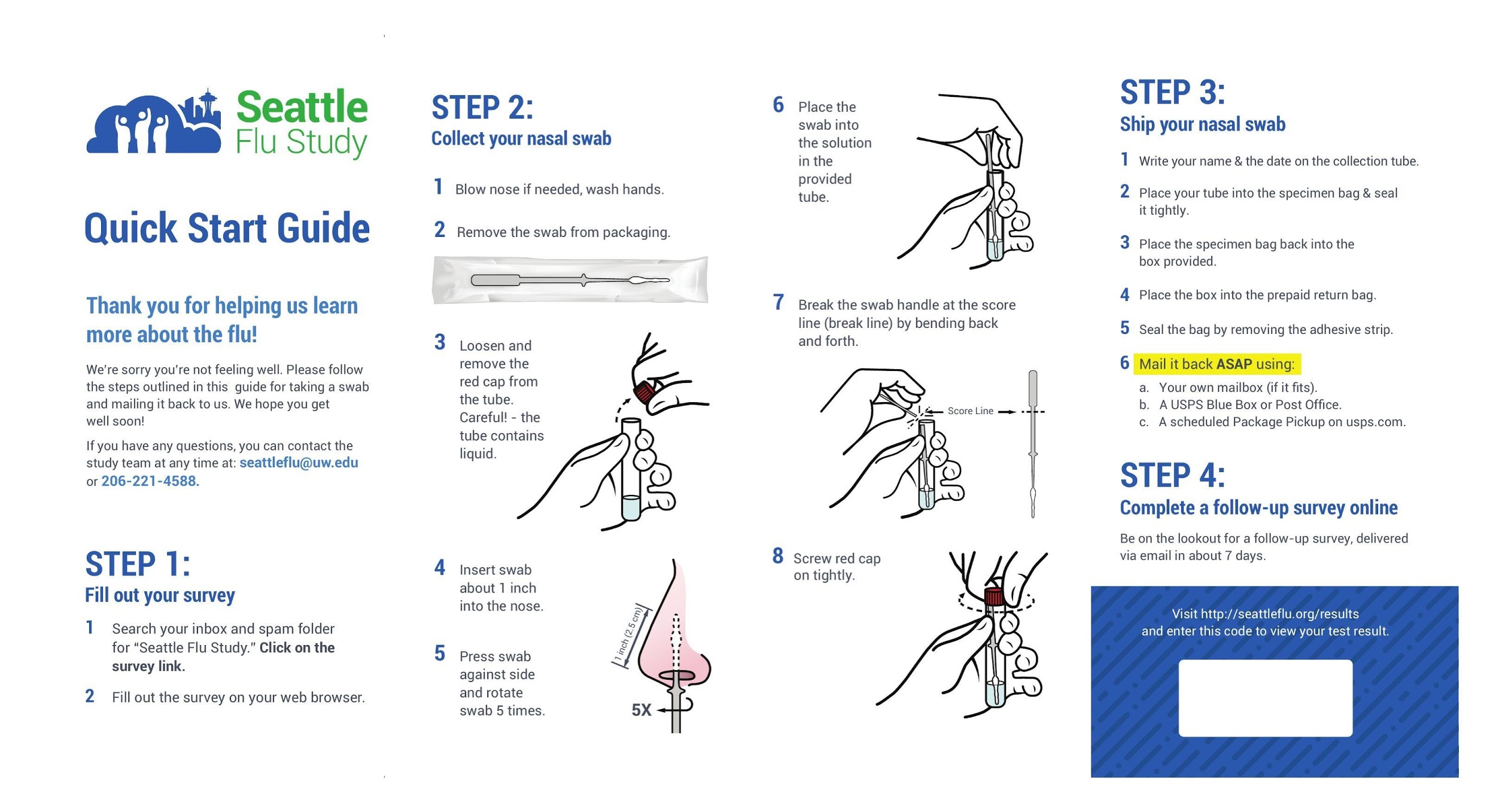
